# CrossPPI: A Cross-Fusion Based Model for Protein–Protein Binding Affinity Prediction

**DOI:** 10.1101/2025.11.25.690371

**Authors:** Suhas Reddy Singam, Nithish Chandra Azad Devarashetty, Sanjana Gogte, Vani Kondaparthi

**Author notes:** Corresponding Author: Vani Kondaparthi.

## Abstract

Many biological processes depend on protein–protein interactions (PPIs), which are particularly important across biology, medicine, and biotechnology. It is essential to accurately predict the binding affinity between protein pairs to prioritize candidate interactions in large-scale studies and expedite drug discovery. The application of cross-attention mechanisms between ligand and receptor protein sequences is often neglected in current computational models, limiting their capacity to accurately represent inter-protein dependencies. In this study, we introduce CrossPPI, a novel deep learning framework that integrates structural and sequential features of interacting proteins to improve binding affinity prediction. To model intricate interactions between protein pairs, CrossPPI uses a transformer-based cross-fusion module and a dual-view feature-extraction approach that combines Graph Attention Networks (GATs) and Convolutional Neural Networks (CNNs).

On the test dataset of 300 protein–protein pairs, CrossPPI achieved a Pearson correlation coefficient (PCC) of 0.7616, a Spearman correlation coefficient (SCC) of 0.7644, a mean absolute error (MAE) of 1.2869, and a root mean square error (RMSE) of 1.6824, indicating its ability to predict the binding affinity of two proteins.

The results highlight CrossPPI’s capability to predict inter-protein binding affinities by leveraging an attention-based integration of sequence and structural features.

## 1. Introduction

Proteins are involved in all significant biological actions, including signalling (1, 2), immunological responses (3, 4), and enzyme-related functions (5, 6), which drive chemical reactions. The most common way proteins enhance their biological activity is through protein-protein interactions (PPIs), which are widely recognized as a major mechanism of altering protein activity. PPIs are a crucial component of protein activity modulation, providing a means for proteins to be functionally active in specific physiological contexts. In many instances, proteins arrive at their biologically active conformational or functional state only through an interaction with a partner protein (7). Therefore, most biological processes rely on PPIs, and PPIs failing or being disrupted in any way can result in loss of biological function, unchecked signaling, and various other negative downstream consequences; PPIs and their dysfunction are associated with many neurodegenerative diseases (8) and cancer (9) disorders. There is also an effort to develop therapeutic modalities that specifically target these PPIs (10).

Researchers utilize experimental methods to investigate PPIs, as well as a variety of classical experimental techniques, including but not limited to: yeast two-hybrid screening (11, 12), isothermal titration calorimetry (13, 14), and surface plasmon resonance (15), to study protein– protein interactions (PPIs). While these methods can be valuable and robust, they can also be expensive, slow, and relatively labor-intensive (16). Over time, computational methods have gained significant traction, providing a faster way to estimate the binding affinity of two protein sequences and enabling researchers to prioritize which interactions to test in the laboratory (17). Early methods in this area generally relied on sequence alignments to make predictions, energy scoring functions, or molecular docking (18). One software tool that predicts the type and number of contacts at the protein interface is PRODIGY (19), but as deep learning (20) has become more prevalent, models using these new methodologies have begun to outperform traditional methods. Some conventional models, like GraphDTA (21) and DeepAffinity (22), predict binding affinity using graph-based molecular representations and sequence-derived embeddings respectively. While these models have been successful in predicting protein–small molecule interactions, there is sparse evidence indicating direct applicability for predicting PPI affinity with the above approaches primarily because these were earlier designed to consider small-molecule binding, which may not encompass the complex, larger interface features of PPIs. Despite the advancements made over the years, predicting PPI binding affinity remains a challenging task (23, 24). A major gap in the current literature is the lack of generalizability, as most models are extensively trained on small or narrow datasets and struggle to transfer their performance to proteins from uncharacterized families. Additionally, approaches that solely focus on either protein sequence or structural information often miss the complementary information available through both forms of data.

In response to the challenges in this area of research, we introduce **CrossPPI**, a deep learning model tailored for predicting protein–protein binding affinity. It is derived from DeepRSMA (25) but modified for protein-protein binding affinity prediction rather than RNA-small molecule binding affinity prediction. CrossPPI model uses structural and sequence information from ligand protein and receptor protein pairs. On the structure side, the model uses Graph Attention Networks (GATs) (26) to model the residue-level interactions and topological relationships. Simultaneously, 1D CNNs (27) are used to acquire hierarchical sequence patterns. The different representations are fused via a cross-fusion transformer module, allowing for bidirectional contextual learning from the ligand and receptor over both sequence and structure modalities. CrossPPI, thus, has the potential to integrate more complex interaction dynamics in a more unified and biologically relevant way as it would relate to the binding affinity of the two interacting partners. The fused embeddings are then pooled to predict the binding affinity.

In addition, CrossPPI’s modular structure offers flexibility and maintainability, enabling easy modification or extension of the model for different applications in protein interaction modelling.

## 2. Method

### 2.1. Overview

The CrossPPI framework consists of three different modules, as illustrated in Figure 1: a pKD prediction module, a cross-fusion module, and a protein feature extraction module, which is used for extracting features for both receptors and ligands. The protein feature extraction module uses a 1D CNN block to extract sequential features from protein residues, and a contact map generated via ESM2(650M) to extract spatial features. The cross-fusion module combines receptor and ligand information, and the pKD prediction module then predicts the pKD for the receptor and ligand protein sequences.

The model’s performance was evaluated with PCC (28), SCC (29), MAE, and RMSE (30, 31). High values for PCC and SCC indicate that the model captured the linear and monotonic relationships, respectively. In contrast, lower values for MAE and RMSE indicate greater accuracy in predicting binding affinity.

### 2.2. Protein feature extraction module

#### 2.2.1. Protein graph findings

The protein graph information is collected from the sequence using ESM2 (650M) to generate a contact map. This contact map is then binarized with a threshold of 0.5, indicating that two amino acids are considered to be in contact if their distance is less than 0.5 (represented as 1) and not in contact if their distance is greater than 0.5 (defined as 0). Three GAT layers are applied to the amino acid embedding derived from the protein sequence, followed by one-hot encoding to learn node embeddings and capture intricate relationships between nodes (i.e., protein residues).

Each edge, or connection between nodes, is given an attention coefficient by GAT, which is determined using the formula given below.

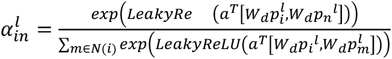

Where LeakyReLU is an activation function (32, 33), *p_i_^l^* is the hidden representation of the amino acid i at the *l*th layer, W_d_ is a trainable parameter and a is the learnable weight matrix, and N(i) is the neighbor nodes of *i*. After acquiring the attention coefficient, the representation of amino acid *i* is calculated by a linear attention aggregation layer:

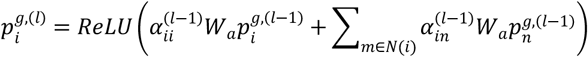

Where W_a_ is a trainable parameter and α is the attention coefficient.

#### 2.2.2. Protein sequence information

After the one-hot encoded sequence has been passed through an embedding layer, the amino acid embeddings were created to extract sequential information from the protein sequence.

Furthermore, the embedding produced was used by ESM2 (650M) to leverage its capacity to capture contextually aware, rich representations of protein sequences (34). The ESM2 (650M) and amino acid embeddings are then both padded to a predetermined length. The embeddings are averaged after padding.

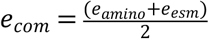

Where e_com_ is the combined embeddings, e_amino_ is the embeddings generated by the embeddings layer, and e_esm_ are the embeddings generated by the ESM2 (650M) model.

To investigate amino acids in various local neighborhoods, three 1D CNN layers with kernel sizes of 7, 11, and 15 were used (35, 36). The final CNN embeddings are obtained by linearly transforming the averaged outputs of the three CNNs.

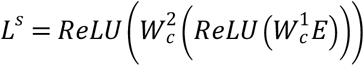

Where E is the averaged embedding, W_c_^1^ and W_c_^2^ are trainable parameters. After averaging the node-level features and masking the padded position, the final sequential embeddings are obtained.

**Fig. 1:**
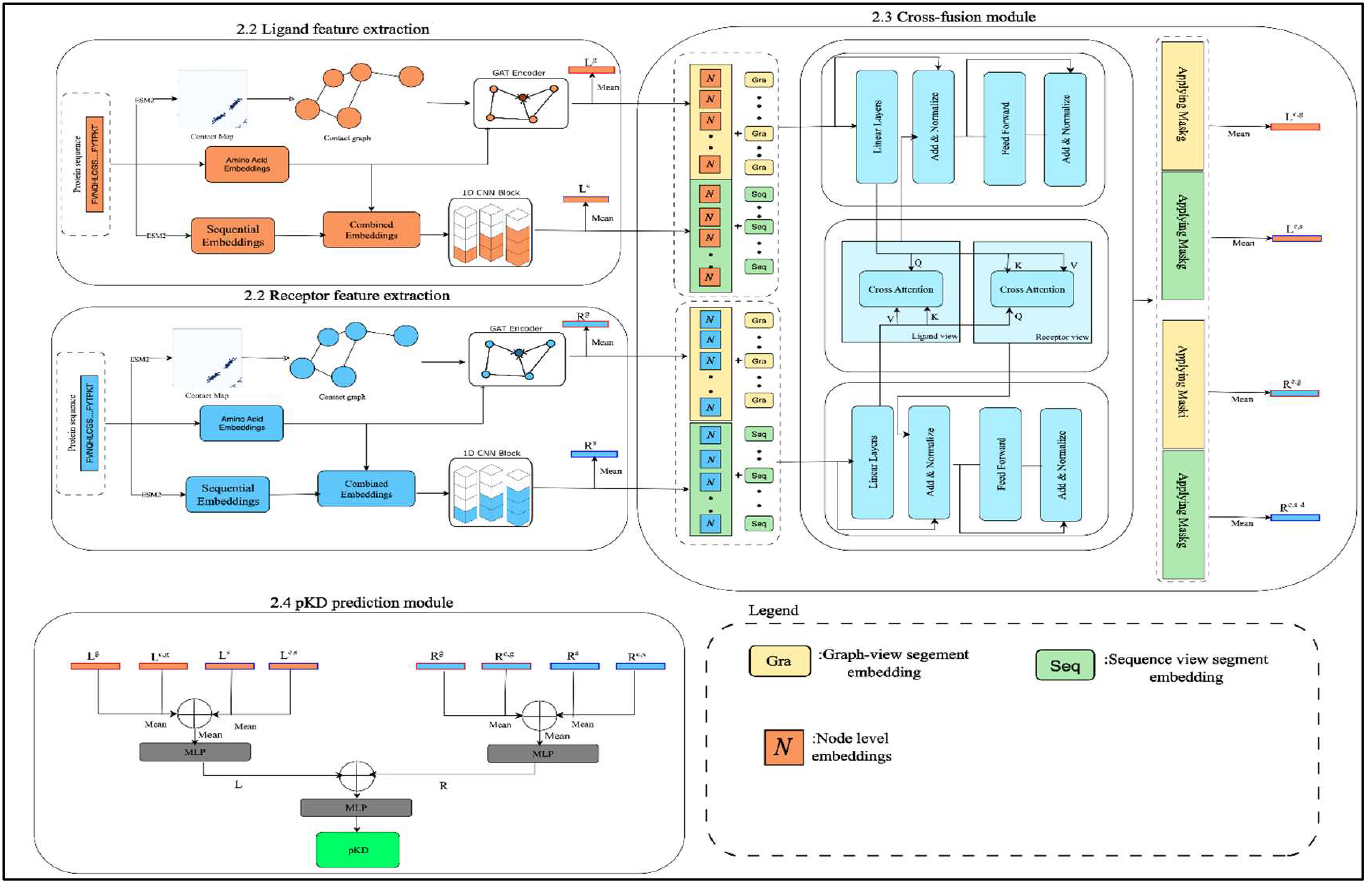
Overview of the CrossPPI framework.

Figure 1 presents 2.2 as the Feature Extraction Module for Ligand and Receptor, as mentioned in the respective section; 2.3 presents the Cross-fusion module to integrate ligand and receptor information from sequential and graphical views, as mentioned. 2.4 represents the Affinity Prediction module to combine the ligand and receptor representations from (2.2) and (2.3) to predict ligand and receptor binding affinity values.

### 2.3. Cross-Fusion Module

To model their interactions for affinity prediction, the cross-fusion module combines the graph and sequence embeddings of the ligand and receptor, produced in sections 2.2.1 and 2.2.2, respectively. To capture intricate, non-local interactions between receptor and ligand residues, it uses a transformer-based cross-attention mechanism.

The two inputs [R^g^, R^s^] and [L^g^, L^s^] that make up the cross-fusion module are the sequential and graphical embeddings of the receptor and ligand, respectively, together with their corresponding masks; to identify the input’s source, it uses a graph-view segment embedding (S^g^) and a sequence-view segment embedding (S^s^). Following the application of cross-attention, the module generates a context layer that includes the fused representations of the Receptor and Ligand, as well as their respective attention ratings. Consider the following representations: r^g^, r^s^, l^g^, and l^s^, which stand for receptor graph, receptor sequence, ligand graph, and ligand sequence embeddings, respectively. Before being concatenated into receptor and ligand hidden states, the graph and sequence view segment embeddings are first added to the receptor and ligand embeddings.

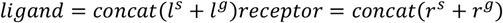

Following this, the masks are concatenated appropriately, the hidden states and masks are grouped, the ligand and receptor hidden states are subjected to cross-attention, and, after masking, SoftMax is applied to the result; the context layers are obtained. Dense layers are applied to the resulting context layers to refine the features; this process is then repeated for n layers to receive the encoded layers and attention scores of the Receptor and Ligand.

### 2.4. Affinity prediction module

The affinity prediction module combines Receptor and Ligand embeddings from Protein feature extraction and cross-fusion modules to predict the binding affinity (pKD). It processes the four Receptor embeddings R^g^, R^s^, R^c,g^, R^c,s,^ and the four Ligand embeddings L^g^, L^s^, L^c,g^, Lc,s .

The Receptor embeddings R and Ligand embedding L are obtained by averaging and then passing their respective embeddings through an MLP. The calculations used to get the embeddings are:

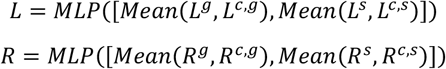

Where the MLP uses ReLU as an activation function and has a dropout of 0.2.

Finally, we use a 3-layer MLP to obtain the predicted pKD of the Receptor and Ligand.

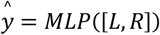

Where, ŷ is the predicted pKD of the Receptor-Ligand pair. The model is trained to minimize the root mean squared error between predicted and actual pKD values. During the training process, the model was trained with a learning rate of 5e-4, hidden dimension of 128, batch size of 32, dropout of 0.2, and weight decay of 1e-7.

### 2.5. Datasets

PPB-Affinity (37) is a protein-protein binding affinity dataset that includes 12062 distinct receptor-ligand pairs, along with their corresponding PDB IDs and experimentally verified KD values in molar units. The dataset examined in this study contains various protein types, allowing for extensive representation of protein-protein interactions. It consists of proteins found in biological organisms, including viruses, bacteria, humans, and non-mammalian organisms. The dataset’s diversity allows the model to learn interactions across species and biological contexts, thereby improving its generalizability and robustness for PPI prediction. For ease of interpretation, the dataset was converted to pKD (pKD = -log10 [KD]). The model was trained on a dataset created by extracting 6436 dimer pairs, that is, only one ligand chain and one receptor chain. Of these, 300 samples were selected from the training set to create a test set for comparative analysis. A density plot, as shown in Figure 2, was designed to visualize the distributions of pKD values in the training and test datasets. The CrossPPI GitHub repository (https://github.com/drugparadigm/CrossPPI) includes all the data and code supporting the article. The dataset related to this study is provided in the accompanying supplementary files (S1 and S2).

**Fig. 2:**
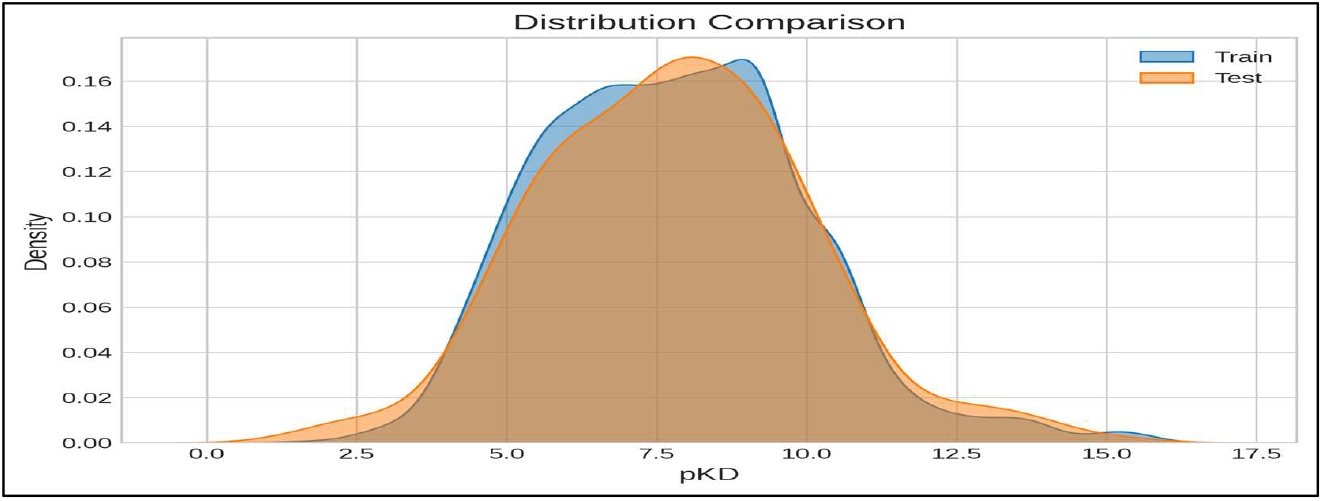
Distribution Curve Comparison of pKD Values.

Figure 2 presents the density distribution curves for pKD values in the train (blue) and test (orange) datasets. The x-axis represents the pKD values ranging from 0.0 to 17.5, while the y-axis indicates the density.

It can be observed from Figure 2 that both the training and test datasets have similar peaks and some overlap, indicating that they have a similar distribution of values.

## 3. Results and Discussion

### 3.1. Five-Fold evaluation

The model’s performance was evaluated using five-fold cross-validation (38) on the 6136-sample training dataset. The dataset is split into k equal parts (in the present case, 5) for cross-validation. The remaining portions are used to train the model, while one of these sections, which comprises 20% of the dataset, is selected as the test set. Each component is used as the test set once during the five repetitions of this procedure. To assess the model’s performance, the PCC, SCC, MAE, and RMSE were computed for each fold. By averaging the PCC, SCC, MAE, and RMSE of the best model from each of the five folds, the final PCC, SCC, MAE, and RMSE of the model were determined, which were 0.7534, 0.7528, 1.1572, and 1.5450, respectively. Table 1 displays the individual folds’ PCC, SCC, MAE, and RMSE values. The best values for each metric are indicated in bold. Additionally, Figure 3 shows the CrossPPI metrics on the training dataset.

**Fig. 3:**
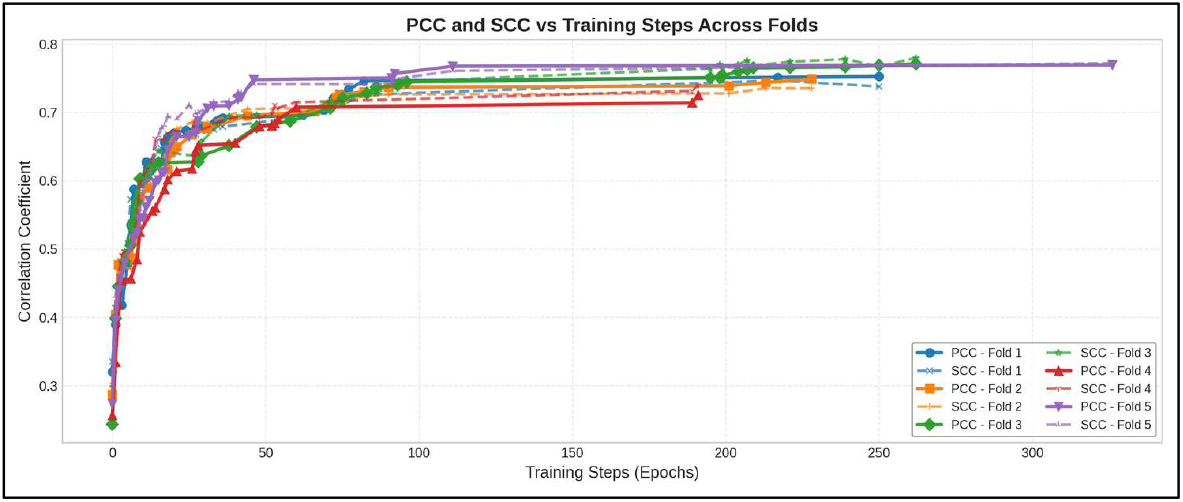
Performance Metrics of CrossPPI on Training Dataset.

The line graph in Figure 3 displays the performance of CrossPPI training metrics (PCC and SCC) over epochs on the training dataset.

**Table 1:** Metrics across all 5-folds.

| Fold | PCC | SCC | MAE | RMSE |
| --- | --- | --- | --- | --- |
| 1 | 0.7526 | 0.7377 | 1.1138 | 1.5154 |
| 2 | 0.7486 | 0.7351 | 1.2033 | 1.5869 |
| 3 | **0.7710** | **0.7796** | **1.0283** | **1.3983** |
| 4 | 0.7253 | 0.7397 | 1.1809 | 1.6013 |
| 5 | 0.7695 | 0.7720 | 1.2598 | 1.6232 |

The metrics in Table 1 present the valid score of each metric across all 5-folds, with the best scores highlighted in bold.

### 3.2. Comparison with other models on the test dataset

The 300-sample test dataset was used to compare the CrossPPI model’s performance to that of other models. The best outcomes are highlighted in bold in Table 2, which presents the results. The CrossPPI model is evaluated with KNN (39) and SVM (40) as the machine learning baselines and transformer (41) and GCN (42) as deep learning baselines. The CrossPPI model obtained a PCC of 0.7616, SCC of 0.7644, MAE of 1.2869, and RMSE of 1.6824 on the test dataset. The test dataset was evaluated using the CrossPPI fold-1 checkpoint. The CrossPPI model’s predictions on the test dataset are displayed as a scatter plot in Figure 4.

**Fig. 4:**
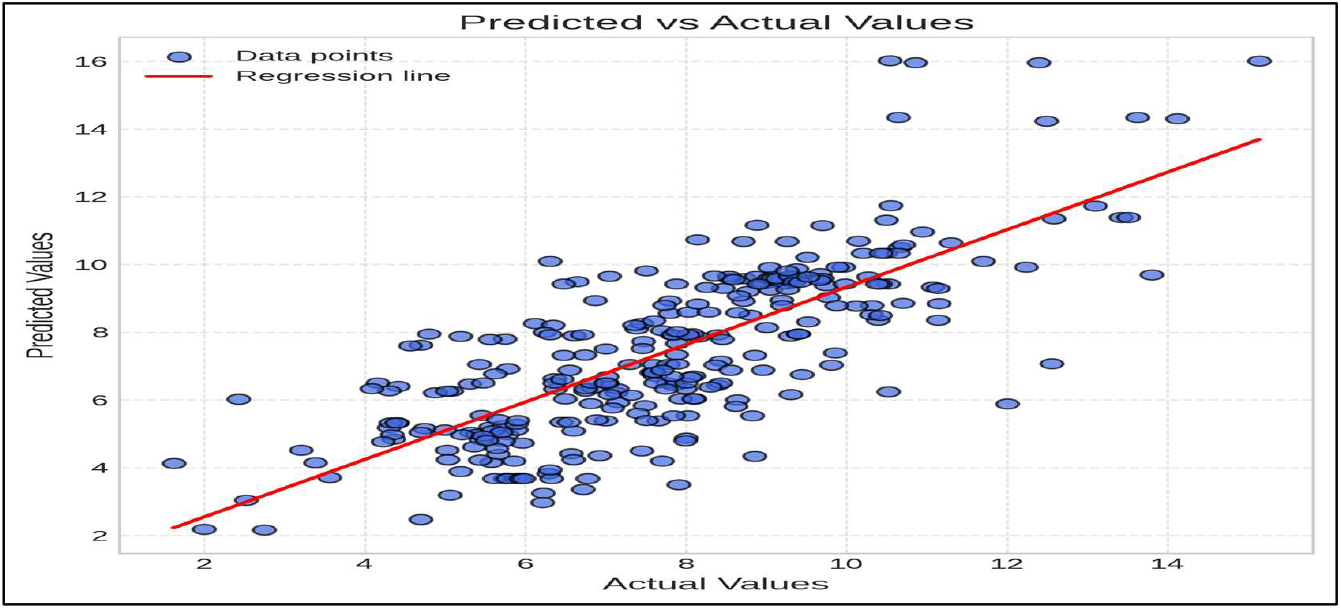
CrossPPI model’s pKD values prediction on the test dataset.

The scatter plot in Figure 4 illustrates the relationship between predicted and actual values, with each blue dot representing a single prediction instance. Points closer to the red line (linear regression line) reflect higher prediction accuracy.

**Table 2:**
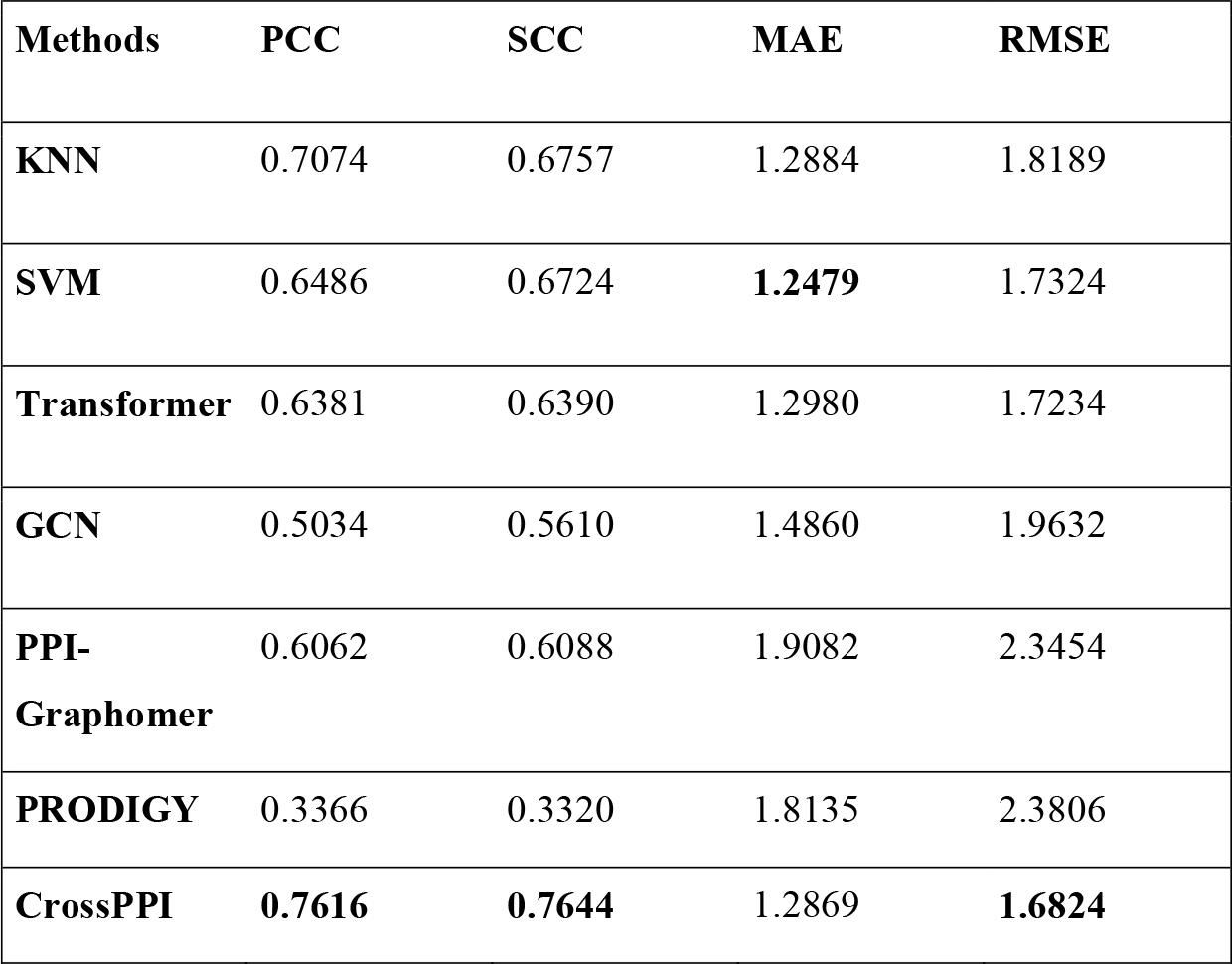
Comparison of models on the test dataset.

All four parameters were compared across all available models and presented in Table 2.

CrossPPI achieved the best PCC, SCC, and RMSE scores among all the models tested and has the second-best MAE.

### 3.3. Ablation Study

To evaluate the contribution of each module in our model, we created three variants and tested their performance using five-fold cross-validation. The three models are CrossPPI without graph context (w/o Graph), CrossPPI without sequence context (w/o sequence), and CrossPPI without cross attention (w/o cross_attn).

The results of the five-fold cross-validation, conducted over 100 epochs for each fold, are shown below in Table 3, with the best outcomes highlighted in bold. The noticeable decrease in the performance of CrossPPI (w/o sequence) highlights the importance of sequence information in accurately predicting protein-protein binding affinity. Additionally, the inferior performance of CrossPPI (w/o cross attention) demonstrates the importance of the model learning the binding patterns of the ligand and receptor sequences.

**Table 3:** Ablation study.

| Method | PCC | SCC | MAE | RMSE |
| --- | --- | --- | --- | --- |
| w/o graph | 0.7442 | 0.7527 | 1.2044 | 1.5413 |
| w/o sequence | 0.7241 | 0.7165 | 1.1747 | 1.5140 |
| w/o cross_attn | 0.7215 | 0.7312 | 1.4084 | 1.7588 |
| CrossPPI | **0.7574** | **0.7584** | **1.1453** | **1.4831** |

Table 3 presents the metrics of ablation studies. Where, w/o means without, and the best performance for each metric is marked in bold.

## 4. Conclusion

The present work, CrossPPI, is a novel deep learning model for predicting protein–protein binding affinity (pKD values). CrossPPI integrates both sequence-based and graph-based representations of protein structures, enabling the model to capture both sequential and spatial information. A primary advantage of the model is its cross-fusion mechanism, which supports effective interaction modeling when the receptor and ligand proteins interact. CrossPPI proposes a new paradigm for pKD value prediction that utilizes complementary sequence and structural features, as well as interaction-aware fusion. Furthermore, because the model uses embeddings and contact maps from the ESM-2 protein language model, improvements to ESM-2 expected in the future will likely enhance CrossPPI’s predictive capability. In addition to its strong predictive accuracy, the model’s modular design allows for easy integration with emerging protein representation techniques and datasets. This flexibility makes CrossPPI a promising foundation for downstream tasks such as PPI site prediction, protein design, and virtual screening in drug discovery. Although our current work focuses on pairwise protein binding affinity, the underlying architecture can be extended to multi-protein systems or protein–small molecule interactions. Future work may also explore integrating experimental data, attention visualization, and interpretability mechanisms to understand the learned interaction features better and guide experimental validation.

## ABBREVIATIONS

CNN: Convolutional Neural Network
ESM2: Evolutionary Scale Modeling 2
GAT: Graph Attention Network
GCN: Graph Convolutional Network
KNN: K-Nearest Neighbors
MAE: Mean Absolute Error
MLP: Multi-Layer Perceptron
PCC: Pearson Correlation Coefficient
PDB: Protein Data Bank
pKD: Negative logarithm of the equilibrium dissociation constant (K_D)
PPIs: Protein-Protein Interactions
PRODIGY: PROtein binDIng enerGY prediction
RMSE: Root Mean Square Error
SCC: Spearman Correlation Coefficient
SVM: Support Vector Machine

## DECLARATIONS

## Availability of Data and Materials

The dataset supporting the conclusions of this article is available in the Zenodo repository https://zenodo.org/records/14271435, and the DOI of that dataset is https://doi.org/10.1038/s41597-024-03997-4.

The source code used for analysis is publicly available at GitHub repository https://github.com/drugparadigm/CrossPPI.

## Supplementary Material

The additional datasets related material are available as supplementary files S1 and S2.

## Ethics approval and consent to participate

The present work did not involve any human participants or animal experiments. Hence, ethical approval was not applicable.

## Author Contributions

S.R.S.: Investigation; validation; methodology; visualization; writing-original draft; formal analysis. N.C.A.D.: Formal analysis; methodology; validation; visualization. S.G.: Conceptualization; methodology; validation; visualization; formal analysis; data curation; software; supervision; writing - review and editing. V.K.: Conceptualization; methodology; writing - review and editing; supervision; formal analysis.

## Consent for publication

Not Applicable

## Acknowledgements

We, the authors SRS, NCAD, SG, and VK, express our sincere gratitude to the Drugparadigm Research Lab for providing the necessary facilities and infrastructure that enabled the successful completion of this work.

